# The genome assembly of the westslope cutthroat trout, *Oncorhynchus lewisi*, reveals interspecific chromosomal rearrangements with the rainbow trout

**DOI:** 10.1101/2024.11.15.623813

**Authors:** Anne-Marie Flores, Kris A. Christensen, Theresa Godin, Yniv Palti, Matthew R. Campbell, Geoffrey C. Waldbieser, Sheron A. Simpson, Brian E. Scheffler, Seth R. Smith, Andrew R. Whiteley, Ryan P. Kovach, Gordon Luikart, Matthew C. Boyer, Marty Kardos, Scott Relyea, Craig Wells, Ben F. Koop

**Author notes:** Corresponding authors (KC), (BK). These authors contributed equally to this work (Anne-Marie Flores, Kris A. Christensen).

## Abstract

Cutthroat trout (*Oncorhynchus clarkii*) are popular among anglers throughout their native range along the West Coast and interior of North America. As they colonized the interior of North America, cutthroat trout diverged into several genetically distinct groups. Many of these groups are now threatened by habitat destruction, hybridization with rainbow trout (*O. mykiss*), and competition from introduced species. These groups were previously classified as subspecies, but recent research suggests they may represent distinct species. In this study, we produced a chromosomal-level genome assembly and a genetic map for one of the species in the cutthroat trout species complex, the westslope cutthroat trout (*O. lewisi*—formerly *O. clarki lewisi*). We also constructed haplotype-resolved assemblies from a westslope cutthroat-rainbow trout F1 hybrid. We used the new genome assemblies to identify major interspecific chromosomal rearrangements between the two sister species, including fusions, fissions, and inversions. These genome assemblies and chromosome data provide valuable insights regarding genetic variation within cutthroat trout and in hybrids between rainbow and cutthroat trout.

**Article Summary:** Westslope cutthroat trout inhabit bodies of water in several states and provinces near the Rocky Mountains and are well-known among anglers. In this study, we produced the first publicly available genome assemblies and a high-density genetic map for this species. These are research tools that allow detailed genetic analyses, including identifying differences among the genomes of related species. These comparisons allow researchers to understand the evolution of these species better and may provide insight into why successful interbreeding is possible among cutthroat and rainbow trout.

## Introduction

Cutthroat trout (*Oncorhynchus clarkii*) are an iconic species of western North America. They hold cultural significance for Indigenous communities and are highly valued for their recreational and economic importance (Mallet and Thurow 2022; DFO 2009). Cutthroat trout are particularly valued by anglers, due to their surface feeding behavior and high catchability (Mallet and Thurow, 2022). A 2015 survey by the Freshwater Fisheries Society of BC (published in the Economic Impact Report - 2019), ranked cutthroat trout as the 4^th^ favourite sport fish among anglers in British Columbia (BC), Canada (gofishbc.com).

Cutthroat trout form a species complex that is native to the coastal and interior waters of western North America (Allendorf and Leary 1988; Behnke 1992; Whiteley et al. 2019). This polytypic species exhibits remarkable phenotypic and genetic variability, diverse life history strategies, and significant evolutionary diversity (Allendorf and Leary 1988; Behnke 2002; Campbell et al. 2011). They occupy various aquatic habitats, including small headwater streams, large rivers, lakes, and estuaries. These habitats share the characteristics of being cold, clean, and oxygen-rich (Behnke 1992; Trotter 2008).

The extensive diversity within cutthroat trout has led to disagreements among researchers regarding its description and classification, particularly concerning the number of species and subspecies (Behnke 1992; Metcalf et al. 2007; Penaluna et al. 2016; Trotter et al. 2018).

Previously, 14 subspecies of cutthroat trout were recognized, two of which are now extinct (Behnke 1992; Behnke 2002). Recent studies (see Wilson and Turner 2009; Metcalf et al. 2012; Loxterman and Keeley 2012; Trotter et al. 2018; Bestgen et al. 2019) have re-evaluated the earlier phylogeny and classification. These studies continue to support the earlier finding that modern cutthroat trout diverged from a common ancestor into four major evolutionary lineages –Coastal, Lahontan Basin, Upper Columbia/Missouri River (Westslope), and the Yellowstone/Southern Rocky Mountain Lineage/Upper Snake River (Figure 1) (Behnke 1992; Allendorf and Leary 1988; Trotter et al. 2018). Furthermore, there may be at least 25 uniquely identifiable evolutionary units (UIEUs) within these lineages (Trotter et al. 2018). The nomenclature and classification of this complex are still subject to debate, and further research is needed to determine the appropriate taxonomic units (species or subspecies).

**Figure 1.**
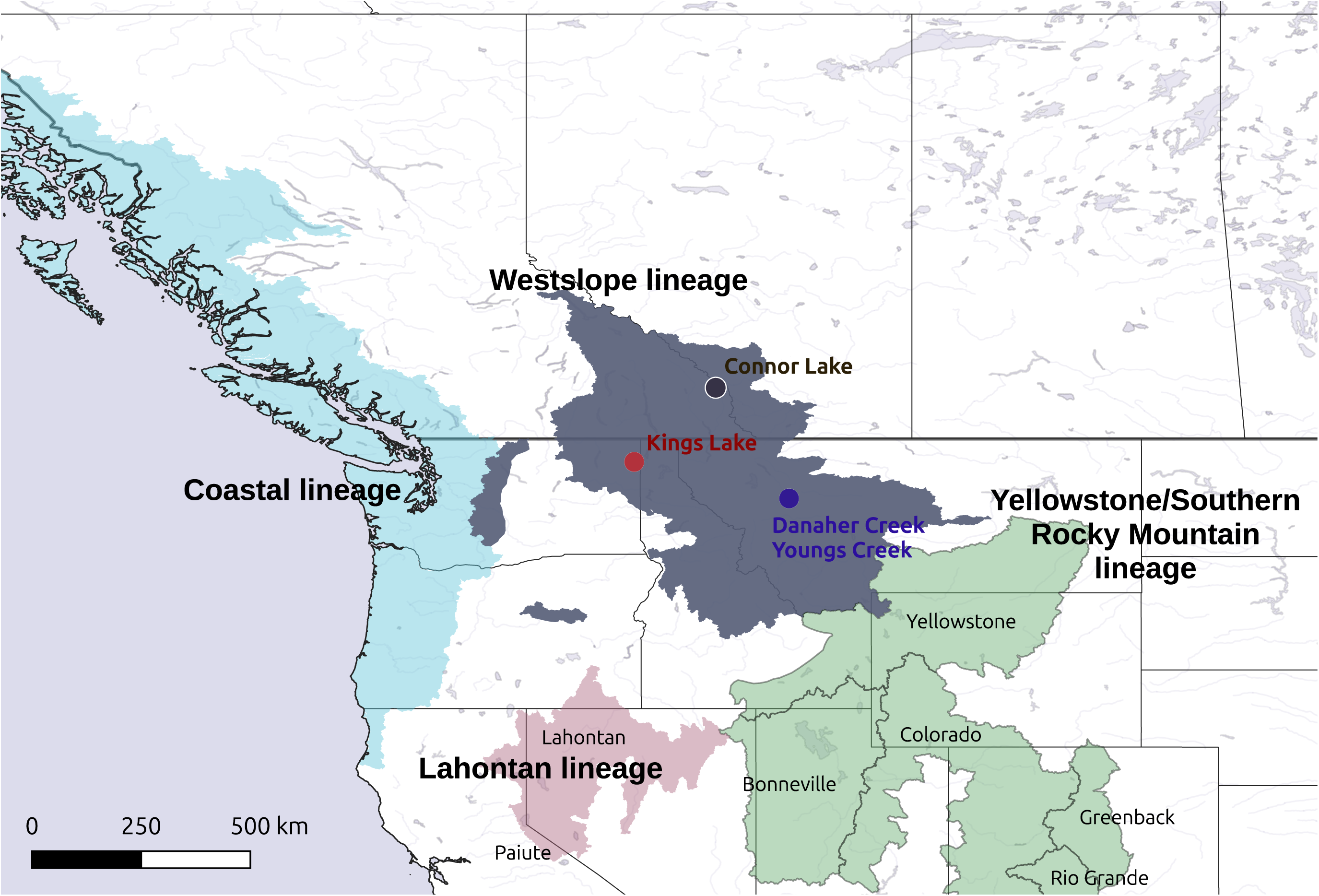
Historical and current distributions of the cutthroat trout in western North America. The four major evolutionary lineages of cutthroat trout are labelled on the map. Westslope cutthroat trout (WCT) were sampled from Connor Lake, British Columbia (Canada), Kings Lake, Idaho (USA), and Youngs Creek and Danaher Creeks in the headwaters of the South Fork of the Flathead River, Montana (USA). Map adapted from Penaluna et al. (2016).

The westslope cutthroat trout (WCT), *O. lewisi*—formerly *O. clarki lewisi* (Page et al. 2023)—is one of the four major evolutionary lineages of cutthroat trout (Figure 1), renowned for its ecological, economic, and cultural significance (Cosewic 2006; Quinn 2005). While the historical distribution of WCT is somewhat uncertain, their range included portions of the Fraser, South Saskatchewan, Missouri, and Columbia River basins (Behnke 1992; Shepard et al. 2005; McPhail 2007). Although these four major basins are currently hydrologically isolated, some degree of historical connectivity or migration among them has been proposed (Young et al. 2018). The WCT lineage can be distinguished from other lineages of cutthroat trout through differences in morphology (Behnke 1992), karyotype (Gold et al. 1977; Loudenslager and Thorgaard 1979) and genomic divergence (Leary et al. 1987; Allendorf and Leary 1988; Utter and Allendorf 1994).

Molecular studies have revealed divergence within the WCT lineage (Drinan et al. 2011; Loxterman and Keeley 2012). Young et al. (2018) proposed that the WCT lineage has differentiated into at least nine UIEUs (Table 1). The phylogeographic structure of WCT across its entire range aligns with major watershed boundaries and cycles of the Pleistocene glaciation (Drinan et al. 2011; Young et al. 2018). Further divergence within the WCT lineage may exist at smaller spatial scales. For example, populations within the neoboreal clade in the Upper Kootenay, Elk, Columbia River, and Fraser Rivers could potentially be classified as separate UIEUs, as they are genetically distinct and associated with major drainage basins in BC (B.C. Ministry of Environment 2014; Taylor et al. 2003).

**Table 1.** List of the proposed uniquely identifiable evolutionary units (UIEUs) that have differentiated from the westslope (Upper Columbia-Missouri River) evolutionary lineage of cutthroat trout. The names primarily correspond to the major river basins. Adapted from Young et al. 2018.

| Name | Location |
| --- | --- |
| John Day cutthroat trout | Upper portion of the John Day River basin |
| Coeur d’Alene cutthroat trout | Upper portion of the Coeur d’Alene River basin |
| St. Joe cutthroat trout | Upper portion of the St. Joe River basin |
| North Fork Clearwater cutthroat trout | Upper portion of the North Fork Clearwater River |
| Salmon cutthroat trout | Salmon River basin |
| Clearwater Headwaters cutthroat trout | Headwaters of the Selway and South Fork Clearwater Rivers |
| Clearwater-Eastern Cascades cutthroat trout | Lochsa River, lower Selway River and lower portion of tributaries to the Middle Fork Clearwater River, and the Wenatchee River, Lake Chelan, and Methow River basins |
| Neoboreal cutthroat trout | Columbia River basin from the Sanpoil River upstream (excluding the Spokane River), Fraser River basin and the South Saskatchewan River basin |
| Missouri River cutthroat trout | Missouri River basin |

Modern populations and the distribution of WCT have significantly declined due to several factors, including habitat loss and fragmentation, barriers to migration, overexploitation, competition from introduced species, climate change, and introgressive hybridization (Cosewic 2006; Shepard et al. 2005; Trotter 2008; Muhlfeld et al. 2014; Kovach et al. 2022). Among these, introgressive hybridization poses the primary threat to the genetic integrity of native WCT populations (Allendorf and Leary 1988; Behnke 1992; Shepard et al. 2005). Hybridization between cutthroat trout, including the WCT lineage, and introduced rainbow trout (RT, *O. mykiss*), is widespread in both Canada and the United States (Young 1995; Rubidge et al. 2001; Boyer et al. 2008; Bennett and Kershner 2009; Rasmussen et al. 2010). In Canada, populations of WCT have been reduced by approximately 80% (Cosewic 2006).

Rainbow and cutthroat trout are sister species (Crespi and Fulton 2004; Wilson and Turner 2009; Crête-Lafrenière et al. 2012), sharing a common ancestor approximately 10.2 million years ago (Stearley and Smith 2016). Despite differences in their karyotypes, Robertsonian rearrangements have maintained the same number of diploid chromosome arms (n = 104) in both species (Loudenslager and Thorgaard 1979; Thorgaard 1983). This consistency in chromosome arm number may contribute to their ability to produce viable, fertile hybrid offspring. Diploid chromosome numbers in cutthroat trout vary from 64 to 68, while in RT, they can vary from 58 to 64 (Loudenslager and Thorgaard 1979; Thorgaard 1983). Hybridization occurs due to the absence of pre-zygotic reproductive barriers and temporal and spatial overlap in spawning (Allendorf and Leary 1988; Behnke 1992).

Introgression between introduced RT and native WCT can lead to the loss of locally adapted traits (Muhlfeld et al. 2009a), reduced fitness (Muhlfeld et al. 2009b; Kovach et al. 2016a), and decreased resilience to environmental changes (Yau and Taylor 2013; Muhlfeld et al. 2014; Kovach et al. 2016b), all of which threaten the survival of WCT populations (Allendorf and Leary 1988; Allendorf et al. 2001; Boyer et al. 2008). Additionally, the introduction of hatchery strains of WCT may affect both the phylogenetic interpretation and the evolutionary integrity of wild WCT populations (Young et al. 2018).

The RT genome assembly was the first published salmonid assembly (Berthelot et al. 2014). Since then, several high-quality assemblies have been produced for various RT strains (e.g., Pearse et al. 2019; Gao et al. 2021). In the absence of a genome assembly, cutthroat trout genomic research has relied on reference-free methods or alignment of sequencing data to divergent reference genomes. To facilitate modern genetic and genomics techniques, we produced a genome assembly of a WCT from a BC lake commonly used as a source for broodstock. We also produced haplotype-resolved assemblies of a WCTxRT F1 hybrid. Hi-C data from the hybrid and a high-density WCT genetic map were used to confirm chromosomal fusions, fissions, and inversions among RT and cutthroat trout assemblies. This study provides the groundwork for research evaluating the genetic basis for reproductive isolation between RT and cutthroat trout, and local adaptation within cutthroat trout lineages.

## Methods

### Sample collection of non-hybridized WCT

Several bodies of water in BC serve as sources of WCT broodstock, with most stocked WCT in BC originating from Connor Lake in the upper Elk System since the 1970s (B.C. Ministry of Environment 2014). Connor Lake, originally barren, was stocked in 1950 with WCT from Kiakho Lake, which itself had been stocked earlier with WCT from various BC sources, including Munroe Lake and Peavine Creek. Although the exact origin of Connor Lake’s WCT is unclear, previous genetic testing has confirmed the trout in Connor Lake are non-hybridized WCT (B.C. Ministry of Environment. 2014). While these trout were once thought to belong to the Yellowstone cutthroat trout lineage (Stenton 1960), genetic and chromosomal analyses have shown that only coastal and westslope lineages exist in Canada (McPhail 2007).

In June 2023, milt samples were collected from three male wild WCT at Connor Lake (Figure 1) by Freshwater Fisheries Society of British Columbia (FFSBC). Provincial authorization to transfer samples was given to FFSBC by Fisheries and Oceans Canada (Licence Number: 134373). Liver tissue was dissected from female cutthroat trout that did not survive egg retrieval. Liver samples were stored in ethanol on ice until they could be transferred to a freezer where they were stored at -80°C before processing.

### DNA extraction of WCT and Oxford Nanopore Technologies (ONT) sequencing

High molecular weight genomic DNA was extracted from the milt using the Nanobind CBB kit (PacBio) according to the manufacturer’s protocol. The Short Read Eliminator Kit (PacBio) was used to reduce the number of small DNA fragments according to the manufacturer’s instructions. Sequencing libraries were then prepared using the Ligation Sequencing Kit V14 (SQK-LSK114, ONT) and sequenced on a R10.4.1 flow cell (FLO- PRO114M) of a PromethION 2 Solo system (ONT). Sequences were generated in FASTQ format using the MinKNOW software (ONT).

### Hi-C library and whole genome shotgun sequencing of WCT

A Hi-C library was prepared from female WCT liver tissue at Canada’s Michael Smith Genome Sciences Centre in Vancouver, BC, Canada. The library was prepared with the Arima High Coverage Hi-C kit (Arima Genomics) according to the manufacturer’s instructions and outlined in the User Guide for Animal Tissues Doc A160162 v01. The resulting proximally- ligated DNA served as the basis for library construction, which also used the NEBNext Ultra II DNA Library Prep Kit (New England Biolabs), Q5 PCR Master Mix, and custom IDT indexed primer pairs. Subsequent library products were amplified with 10 reaction cycles using NEBNext Q5 Master Mix (New England Biolabs) supplemented with 2 mM MgSO_4_. The amplified products were bead cleaned, quantified, and then sequenced on the Illumina NovaSeq platform (Illumina) using PE150 sequencing.

Genome shotgun sequencing was also performed at Canada’s Michael Smith Genome Sciences Centre using the TruSeq DNA PCR-free library prep kit (Illumina) automated via a Microlab NIMBUS liquid handling robot (Hamilton Robotics). This method was chosen to reduce bias and coverage gaps from PCR amplification in high GC or AT-rich regions. Briefly, 500 ng of genomic DNA (extracted from milt of a different male from ONT sequencing – due to limited DNA) was placed into a 96-well microtitre plate and fragmented using sonication (Covaris LE220). The fragmented DNA was end-repaired and size-selected using paramagnetic PCRClean DX beads (C-1003-450, Aline Biosciences), which targeted a 300-400 bp range. After 3’ A-tailing, full-length TruSeq adapters were added. The libraries were purified with paramagnetic beads from Aline Biosciences and quantified using a qPCR Library Quantification kit (KAPA, KK4824). The libraries were then sequenced with paired-end 150 bp reads on an S4 (v1.0) flowcell of the Illumina NovaSeq 6000 sequencer following the manufacturer’s protocol.

### Genome assembly - WCT

Flye (version 2.9.2) (Kolmogorov et al. 2019) was used to produce the initial WCT contig genome assembly from the ONT reads. The --nano-hq, --read-error 0.03 -g 2.5g --asm-coverage 40 parameters were used for this initial assembly. This assembly was then polished with racon (version 1.5.0) (Vaser et al. 2017) once and Pilon twice (version 1.24) (Walker et al. 2014).

Minimap2 (version 2.17) (Li 2018; Li 2021) was used to align the ONT reads to the initial assembly (parameters: -ax map-ont) before the alignment was polished using racon (parameters: -u). Short-reads were used for polishing using Pilon after they were trimmed using Trimmomatic (version 0.39) (Bolger et al. 2014) with the following parameters: TruSeq3- PE.fa:2:30:10:2:keepbothReads, LEADING:3, TRAILING:3, MINLEN:36. The trimmed reads were aligned to the racon polished assembly with bwa mem (version 0.7.10) (Li and Durbin 2009; Li and Durbin 2010; Li 2013) with default settings. After the alignments were sorted with SAMtools (version 1.9) (Li et al. 2009; Danecek et al. 2021), default Pilon parameters were used to polish the assembly.

After polishing, the Arima Genomics mapping_pipeline (version 3, see github.com) was used to map the Hi-C data to the assembly. The alignments were sorted using SAMtools, and Matlock and Juicebox utilities (Phase Genomics, see github.com) were used to convert the alignment file to a Hi-C contact map. Juicebox (version 1.11, Durand et al. 2016) was used to visualize the Hi-C contact map and to make manual changes to the order and orientation.

Alignments of the assembly to the Swanson rainbow trout genome assembly (version 2, GCA_025558465.1) were produced using Dgenies (Cabanettes and Klopp 2018) and were used to verify order and orientation of scaffolds. Juicebox utilities were used to output the modified assembly.

### Sample collection and production of F1 hybrid WCTxRT

A cross was generated between a female WCT (Kings Lake strain) from the Cabinet Gorge Fish Hatchery, Idaho and a coastal male RT (*O. m. irideus,* Hayspur strain) at the Hayspur Hatchery, Idaho, USA in May 2022 (Figure 1). Fin tissue samples were collected from the sire and dame of the cross. Blood was sampled from a single male F1 hybrid offspring when it was approximately one year in age. The Kings Lake strain was tested for introgression with RT, and RT alleles were identified at a frequency of less than 1% (Idaho Fish and Game, unpublished data).

### Haplotype-resolved genome assembly – F1 hybrid WCTxRT

DNA was extracted from a blood sample of a single F1 hybrid male offspring (NCBI BioSample accession SAMN43529089 and BioProject PRJNA1157974). HiFi reads were produced using the CCS mode on the PacBio Sequel II system (GBRU Stoneville, MS). A total of 117.5 Gb sequence data (49x) was generated from 7.6 million HiFi reads (NCBI SRA accessions SRX26069220 and SRX26069221). The Hi-C libraries were prepared from a frozen blood sample and sequenced by a commercial vendor (Phase Genomics, Seattle, WA) to produce 868 million Illumina paired-end reads (2x150 bp) (Accessions SRX26061379, SRX26061380, SRX26061381). Illumina whole genome sequencing data was generated by a commercial vendor (Admera Health, South Plainfield, NJ) from the WCT female parent (SAMN43529090) and RT male parent (SAMN43529091). In total, 89.1Gb (37x) and 94.4Gb (39x) were generated from the WCT (SRX26028960) and RT (SRX26028961) parents, respectively.

The genome assembler Hifiasm (Cheng et al. 2021) was used to generate two genome-wide haplotypes (WCT and RT) from the F1 hybrid, using input data from PacBio HiFi long read reads, Hi-C Illumina sequences, and whole genome short reads sequence data from both parents. The total length of the RT contigs assembly was 2.33 Gb in 3,489 contig sequences and the WCT contigs assembly was composed of 3,808 sequences for a total combined length of 2.35 Gb. The order and orientation of contigs was inferred manually from Hi-C data (see previous section for full details).

### Restriction site associated DNA sequencing (RADSeq) genetic map

The genetic map was constructed using genotypes of parents and offspring from two haploid families and one diploid family bred at the Sekokini Springs Fish Hatchery operated by Montana Fish, Wildlife, and Parks (MTFWP). Linkage mapping families were spawned using parents collected from Youngs Creek and Danaher Creeks in the headwaters of the South Fork of the Flathead River, Montana, USA (Figure 1). The linkage map was constructed following the methods of Blumstein et al. (2020) and Waples et al. (2016). The linkage map and a detailed description of the linkage mapping methods are available in the Supplementary Data (File S1 and File S2). The WCT genetic map was compared to version one of the Swanson RT genome assembly (version 2 was not available at the time it was created). Version one and two are similar, with the exception of an inversion on Omy05.

### Comparison of genome assemblies and genomic metrics

Dgenies (Cabanettes and Klopp 2018) was used to align the three genome assemblies produced in this study to the second version of the Swanson RT genome assembly (Pearse et al. 2019, GCA_025558465.1). The Arlee (Gao et al. 2021, GCF_013265735.2), Whale Rock (GCA_029834435.1), and Keithly Creek RT (GCA_034753235.1) genome assemblies were also aligned given the substantial genetic variation between RT strains (Palti et al. 2014). We also verified the structural changes of the WCT genome assemblies with the genetic map.

The WCT genome assembly was also compared with a genetic map generated from a Yellowstone cutthroat trout (YCT, *O. clarkii bouvieri*) hybrid (Ostberg et al. 2013). We aligned markers from the YCT genetic map (Ostberg et al. 2013), originally taken from Rexroad et al. (2008), to the WCT genome assembly using BLASTN (Altschul et al. 1990, using the outfmt 6 parameter). The output was filtered for the best alignments and we were able to assign linkage groups from the genetic map to chromosomes in the genome assembly.

Length and contig N50 values were calculated using the index file generated by SAMtools from the genome assemblies. BUSCO (version 5.5.0, Manni et al. 2021) scores were used to access genome completeness. These analyses were conducted on the public Galaxy server (usegalaxy.org; Afgan et al. 2016) using the actinopterygii_odb10 gene set. We also used Mequry (version 1.4, Rhie et al. 2020) to determine completeness (kmer based, k=21) and to determine the estimated error rate. These analyses used trimmed short reads for the comparison (see above). For the haplotype-resolved genome assemblies of the WCTxRT F1 hybrid, we used short-reads from the parents and the progeny to generate hap-mers and identify the completeness of each haplotype-resolved assembly.

### Sex chromosomes

The salmonid sex-determining locus, sdY, is located approximately 5 Mb from the start of Omy29 (a.k.a. OmyY) in the Swanson and Arlee RT genome assemblies (Pearse et al. 2019; Gao et al. 2021). We used BLASTN (Altschul et al. 1990) and the sdY gene sequence from RT (Genbank AB626896.1) to try to locate the scaffold containing sdY in the WCT and the F1 WCTxRT hybrid genome assemblies.

## Results

We generated a chromosome-level reference genome assembly for a WCT and haplotype-resolved genome assemblies from a WCTxRT F1 hybrid (Table 2). The WCT genome assembly was 2.4 Gb in length, with a contig N50 of 5.0 Mb and 98.2% complete BUSCOs (Table 2). The hybrid WCT haplotype-resolved assembly was also 2.4 Gb long and had 98.1% complete BUSCOs. The contig N50 was 2.0 Mb. Similarly, the hybrid RT haplotype-resolved assembly was 2.3 Gb long, with a contig N50 of 2.0 Mb and 98.1% complete BUSCOs.

**Table 2.** Genome sequencing and assembly results for a non-hybridized westslope cutthroat trout (WCT) and haplotype-resolved genome assemblies from a WCT x rainbow (RT) F1 hybrid.

|  | WCT | Hybrid WCT | Hybrid RT |
| --- | --- | --- | --- |
| Total length (Gb) | 2.4 | 2.4 | 2.3 |
| Contig N50 (Mb) | 5.0 | 2.0 | 2.0 |
| Complete BUSCOs | 98.2% (S:55.1, D:43.1) | 98.1% (S:55.6, D:42.5) | 98.1% (S:55.5, 45.6) |
| QV* | 43.42 | 43.62 | 43.63 |
| Error rate | 4.6e-05 | 4.3e-05 | 4.3e-05 |
| Completeness† | 97.1% | Cutthroat: 98.27%<br>Rainbow: 0.43% | Cutthroat: 1.98%<br>Rainbow: 98.75% |

We identified interspecific chromosomal rearrangements between WCT and RT with whole genome assembly alignments and confirmed putative rearrangements using the hybrid Hi-C data (Figures 2 and 3). In contrast to the RT genome assemblies, the WCT genome, the WCT portion of the hybrid, and the WCT genetic map displayed no variation in chromosomal arrangements (Figure 2).

**Figure 2.**
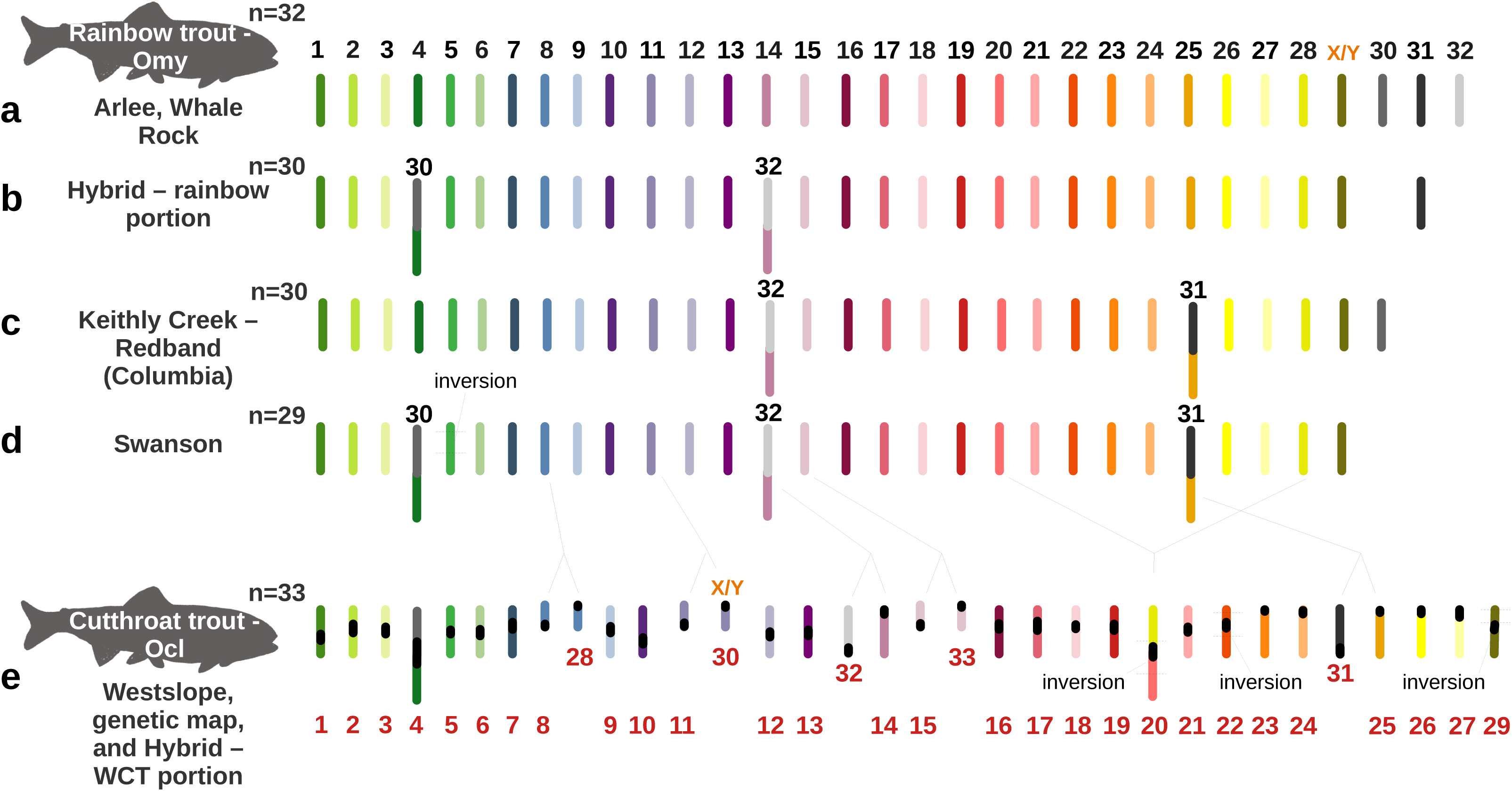
Comparison of syntenic chromosomes among multiple strains of rainbow trout (RT) and westslope cutthroat trout (WCT). This figure highlights karyotype variation among Arlee and Whale Rock RT (a), the RT portion from the WCTxRT hybrid (b), Keithly Creek RT (c), and Swanson RT (d) strains. Only one RT intraspecific inversion was identified on Omy5 (d). The WCT genome assembly (Connor Lake), the WCT genetic map, and the WCT portion of the WCTxRT hybrid all share the same karyotype (e). The potential sex chromosome for WCT is indicated with X/Y. Chromosome numbering is shown in black for RT (top) and red for WCT (bottom). While chromosomes are not to scale, centromeres illustrated on the WCT (black ovals) are depicted relative to their size (based on the genetic map centromere positions). Interspecific inversions, relative to RT, are shown with dashed lines (e). They include pericentric inversions on Ocl20 and Ocl22, and a paracentric inversion on Ocl29.

**Figure 3.**
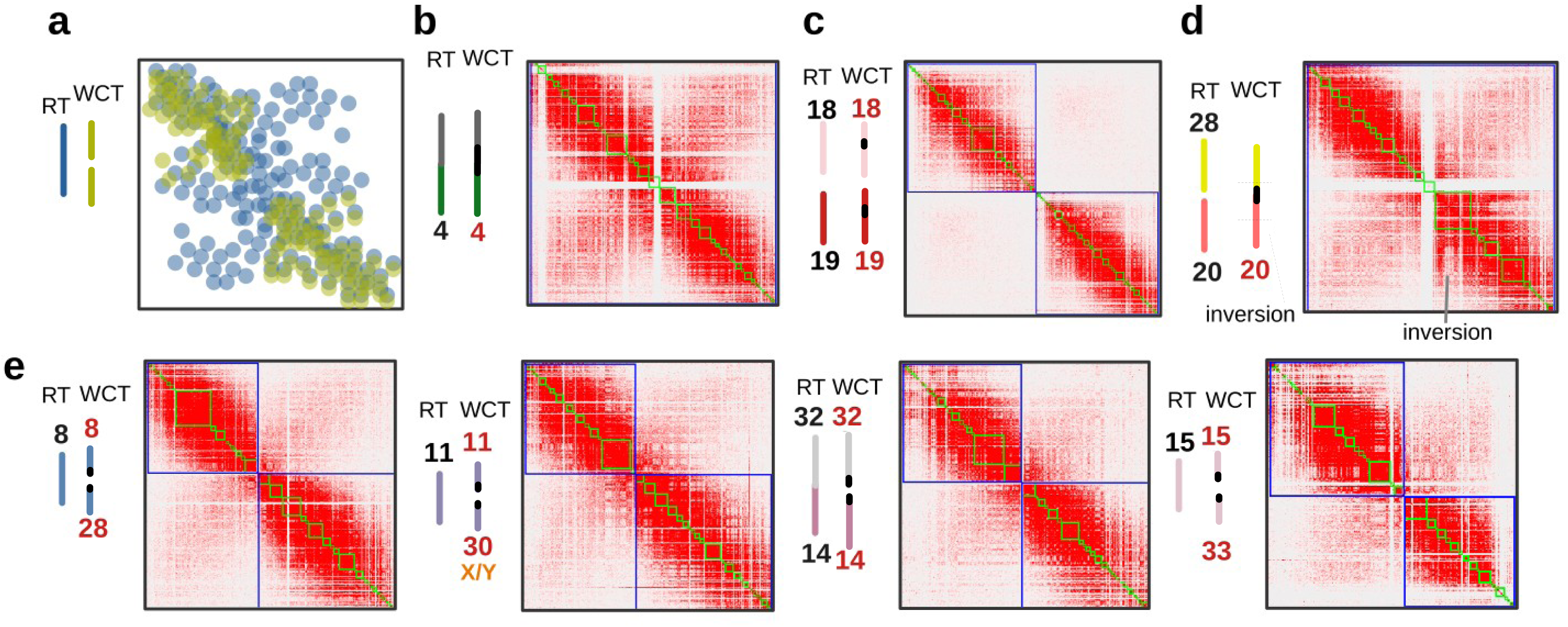
Verification of fusions, fissions and inversions between rainbow trout (RT) and westslope cutthroat trout (WCT) using WCTxRT hybrid Hi-C data. (a) An illustration to demonstrate how Hi-C interaction patterns (right) from two different parental chromosomes (left) are visualized in Juicebox as the hybrid chromosome when the parental chromosomes hypothetically have different colors. In this example, RT is represented by a fused chromosome (blue – bottom layer), and WCT by two unfused chromosomes (green – overlapped layer). The density of the points reflects the degree of interaction between sections of a chromosome (a proxy for physical distance between those sections). Where the RT chromosome is fused, there are fewer interactions in the hybrid overall because of the lack of interactions from the WCT unfused chromosomes. In practice, Juicebox uses a single colour scale, so identifying these features depends entirely on variations of the intensity of the interaction matrix. (b) An example in the hybrid where both parental chromosomes are not polymorphic. Chromosome boundaries are outlined in blue squares (relative to the WCT) and the green boxes are contigs (interior boundaries). (c) An example of two unfused chromosomes in both parental genomes. (d) An example of a fusion in the WCT parental genome and an interspecific inversion. Notice the decreased intensity of the interactions near the center relative to (b). (e) Examples of fissions in the WCT parental genome.

The WCT has 33 chromosomes, compared to the 29 chromosomes observed in the Swanson RT strain, and lacks the double inversion on chromosome 5 (Omy05) found in some RT (Pearse et al. 2019). Five interspecific fissions and one fusion were identified in comparisons of WCT with RT. The WCT also has interspecific pericentric inversions on Ocl20, Ocl22, and a double paracentric inversion on Ocl29. The inversion on Ocl20 occurs in the same region as the Omy20 chromosomal inversion previously characterized in Hale et al. (2023) (Figure 4). Smaller interspecific inversions were also present in the WCT genome, though these were not well supported by the Hi-C data.

**Figure 4.**
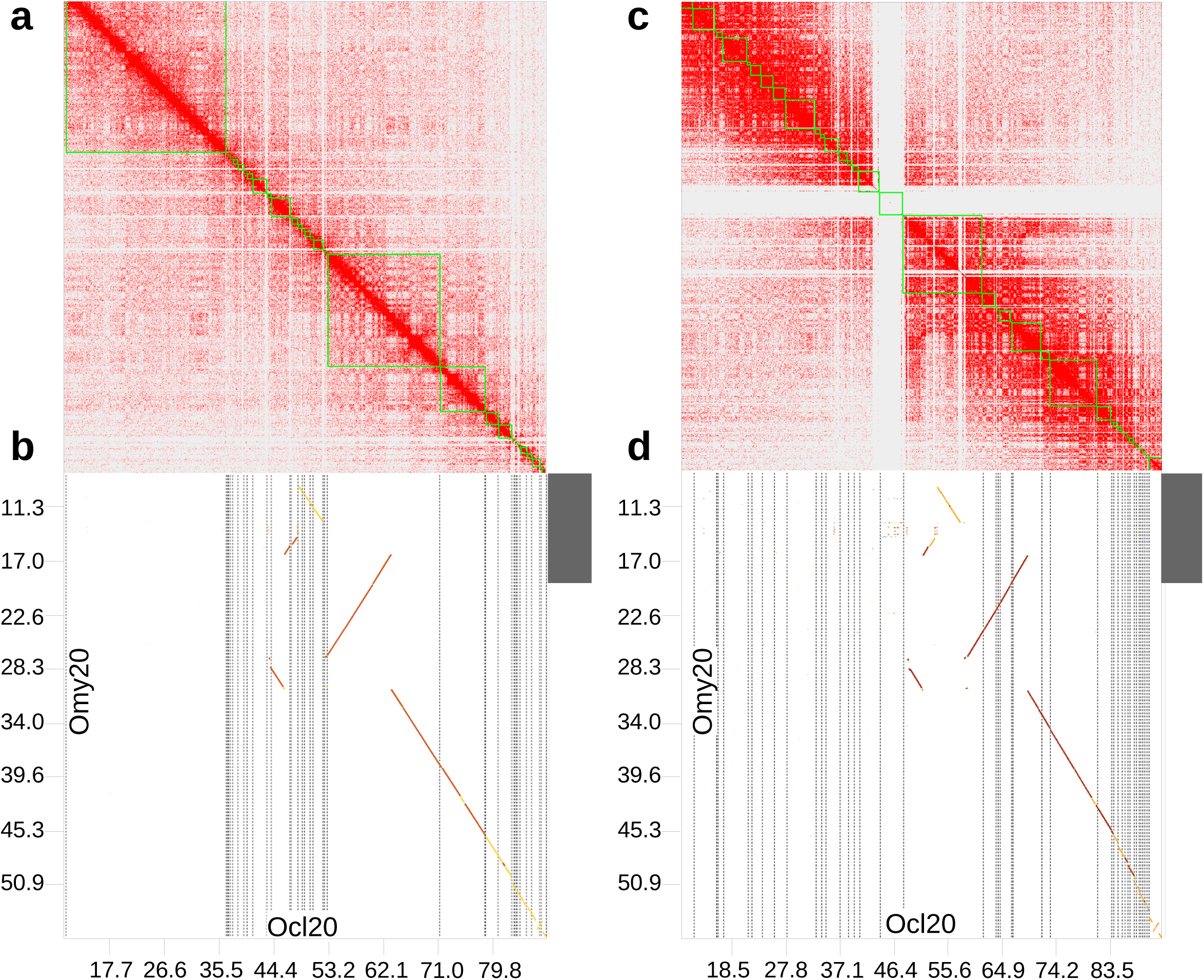
Ocl20 interspecific chromosomal inversion relative to Omy20. (a) Hi-C contact map for part of the WCT Ocl20 (Connor Lake, WCT assembly), with each point representing a contact along the chromosome between locations on the vertical and horizontal axes. Darker points represent more contacts (a proxy for the distance between points, more contacts are expected for locations closer together). Boxes serve as contig boundaries that comprise the Ocl20 chromosome in this region. (b) Dot plot of alignments from contigs in the Connor Lake, WCT assembly to the rainbow trout Omy20 (axis units – Mb). Each vertical line symbolizes a contig boundary and the diagonal lines represent the alignment to Omy20. The gray bar (top right) indicates a part of the boundary from the intraspecific and polymorphic Omy20 chromosomal inversion characterized in Hale et al. (2024). (c) Hi-C contact map of the WCTxRT hybrid. Since the WCTxRT hybrid had parental haploid genomes from both species, the Hi-C contact map from the hybrid will have conflicting data where there are differences between species. Contacts from the WCTxRT hybrid in this region are dense in conflicting locations, suggesting that the WCT and RT chromosomes in the hybrid are different from each other (specifically, the contig within the 46.4-64.9 Mb interval). This supports the rearrangement identified from alignments to the rainbow trout genome assembly. (d) Dot plot of the WCT haplotype-resolved genome assembly (WCT portion of the WCTxRT genome) to Omy20.

For the hybrid RT portion, the genome assembly is similar to the Swanson strain; however, it lacks the Omy05 inversion and the fusion event between chromosomes Omy25 and Omy31 (Gao et al. 2021). The hybrid RT has 30 chromosomes, similar to the Keithly Creek RT genome assembly (Figure 2). The hybrid RT exhibits two notable fusions: chromosomes Omy4 with Omy30 and chromosomes Omy14 with Omy32. The key difference between the hybrid genome assembly and the Keithly Creek genome assembly is the fusion of different chromosome arms, with the hybrid having Omy4 fused with Omy30 instead of Omy25 fused with Omy31.

We located the sdY gene in the WCT genome assembly. Our results suggest that the sex chromosome (X/Y collapsed into one chromosome as this was a haploid assembly) is Ocl30 (which is different from RT). This evidence is weak due to limited Hi-C contact points observed in Juicebox (data not shown). We were unable to find the sdY gene in either of the haplotype-resolved genome assemblies from the F1 hybrid.

## Discussion

The three de novo genome assemblies generated in this study demonstrated high completeness, with BUSCO scores above 98%. The two haplotype-resolved assemblies had lower contig N50 values compared to the WCT assembly. This could be due to differences in sequencing technologies used for the different assemblies, different assembly strategies, or differences in sequencing coverage. The chromosome-level genome assembly of the WCT and the haplotype-resolved assemblies from the WCTxRT F1 hybrid provide valuable insights into the genomic structure and evolutionary dynamics of cutthroat and rainbow trout.

Comparative analysis between the WCT portion of the hybrid genome, the WCT assembly, and the Swanson RT assembly revealed several distinctive interspecific chromosomal rearrangements. The WCT assembly has 33 haploid chromosomes, consistent with earlier karyotypic studies by Loudenslager and Thorgaard (1979) and the WCT genetic map produced in this study. Despite differences in chromosome number, the number of chromosome arms remains constant, indicating that chromosomal variations are primarily due to fusions or fissions (Thorgaard 1983; Phillips and Rab 2001). The higher chromosome number in WCT compared to the Swanson RT results from the fission of Omy08 into Ocl08 and Ocl28, Omy11 into Ocl11 and Ocl30, Omy15 into Ocl15 and Ocl33, Omy14 into Ocl14 and Ocl32 and Omy25 into Ocl25 and Ocl31. The only fusion in WCT relative to RT was the fusion of Omy20 and Omy28 into Ocl20.

The chromosomal rearrangements in WCT closely resemble those reported by Ostberg et al. (2013) in YCT, which belong to the Yellowstone/Southern Rocky Mountain/Upper Snake River evolutionary lineage (Figure 1). In that study, a genetic map that was produced using YCTxRT hybrids identified several chromosomal rearrangements that differed from the RT genome. Here, we compared the genomes of YCT and WCT by mapping the flanking sequences of the microsatellites that were used by Ostberg et al. (2013) to generate the YCTxRT genetic map (Rexroad et al. 2008; Ostberg et al. 2013) onto the WCT genome assembly, and found that the main difference between WCT and YCT is their chromosome number. The YCT genome has 32 haploid chromosomes compared to 33 in the WCT genome (Loudenslager and Thorgaard 1979). This difference is attributed to the absence of a fission on chromosome Ocl08 in YCT.

Chromosomal inversions play a significant role in behavior, local adaptation, and diversification by suppressing recombination, which can drive evolutionary change (Ostberg et al. 2013; Wellenreuther and Bernatchez 2018; Pearse et al. 2019). The WCT genome has three inversions relative to RT: a double pericentric inversion on chromosome Ocl20 (separate inversions in each species that overlap), a pericentric inversion on Ocl22, and a paracentric inversion Ocl29. The 10 Mb inversion on Ocl20 in WCT relative to RT overlaps with the 14 Mb inversion on Omy20 that was previously documented in rainbow trout populations (Pearse et al. 2019; Campbell et al. 2021; Gao et al. 2021; Hale et al. 2024). Although the two inversions overlap, they appear to have originated in separate events as they do not share common start and end positions. Interestingly, a pericentric inversion of the entire q-arm of Omy20 in YCT was hypothesized as one of the rearrangements that occurred before the Omy20-Omy28 fusion in YCT and WCT (Ostberg et al. 2013). Ostberg et al. (2013) suggested that this inversion was necessary for the fusion to occur. In rainbow trout, the Omy20 inversion predates the Omy05 inversion and protein-coding genes found within the inversion are associated with growth, reproduction, immune function, and early development (Cádiz et al. 2021; Hale et al. 2024).

The WCT genome and the hybrid RT genome lack the derived form of the Omy05 inversion previously described in Swanson RT (Pearse et al. 2019; Weinstein et al. 2019). The Omy05 double inversion spans approximately 55-Mb and is associated with adaptive traits such as life-history development (residency vs. anadromy), sexual maturation, and behavior (Sundin et al. 2005; Miller et al. 2012; Pearse et al. 2019; Rundio et al. 2021). The absence of the Omy05 inversion in the two WCT genomes and the genetic map examined in the current study is likely a consequence of the age of the Omy05 inversion, which occurred after the divergence of rainbow and cutthroat trout (Hale et al. 2024).

For the remaining inversions observed in the WCT genome relative to the RT, it is unclear whether they are unique to WCT or shared with the other cutthroat trout lineages. Smaller inversions were identified in the WCT genome relative to the RT, though these were not well supported by Hi-C data. Further research is needed to establish whether these smaller inversions represent true genomic variants or are artifacts of contigs misorientation during genome scaffolding.

Chromosome rearrangements are known to suppress recombination events in rainbow and cutthroat trout hybrids (Ostberg et al. 2013). In this study, up to nine WCT chromosomes would be expected to have recombination suppression in an F1 hybrid between WCT and RBT. While evidence suggests reduced recombination near polymorphic fusion/fission events, we lack data from post-F1 generations to support this. In a North American Atlantic salmon strain, non-Robertsonian translocations showed recombination suppression near fusions, but polymorphic Robertsonian translocations only exhibited this suppression in one out of ten female genetic maps (MacLeod-Bigley and Boulding 2023). If these rearrangements become common following hybridization, Robertsonian translocations may exhibit infrequent recombination suppression, as seen in Atlantic salmon.

## Conclusion

Our study provides new insights into the genomic architecture of cutthroat trout and its divergence from rainbow trout. The unique chromosomal features of WCT, combined with comparisons to previous studies on hybrid trout genomes, enhance our understanding of the evolutionary and adaptive processes shaping these trout species. The three genome assemblies and the genetic map offer a valuable resource for future research into the genetics, ecology, and conservation of cutthroat trout and their hybrids with rainbow trout.

## Data Availability

All WCT sequence data are associated with the NCBI BioProject: PRJNA1151023. The F1 hybrid and parental DNA sequences are associated with BioProject PRJNA1157974. The genetic map is available as supplemental material (Files S1 and S2). Pipelines used in this work and which are not published can be found at https://github.com/ArimaGenomics/mapping_pipeline and https://github.com/phasegenomics/juicebox_scripts.

## Acknowledgments

We thank Dr. Guangtu Gao with our utmost gratitude. Sadly, Dr. Gao passed away last year. Before his passing, he produced the haplotype-resolved genome assemblies of the WCTxRT hybrid. We would like to take this opportunity to acknowledge his incredible talents and his important contribution to this work and to salmonids genomics research.

We thank the Michael Smith Genome Science Centre and their technicians for producing the Hi-C and WGS data. Thanks also to the Digital Research Alliance of Canada (alliancecan.ca) and its regional partner at the University of Victoria (BC DRI Group) for computational resources. We appreciate the extensive help in sample collection that the Freshwater Fisheries Society of BC provided. We thank Vibha Tripathi for her assistance with sequence data submission to NCBI-SRA. We thank Jim Seeb and the Sekokini Hatchery for their help making crosses. We also thank Brooke Penaluna for sharing GIS files to produce the map.

Mention of trade names or commercial products in this publication is solely for the purpose of providing specific information and does not imply recommendation or endorsement by the U.S. government. USDA is an equal opportunity provider and employer.

## Supporting Information

**File S1. Linkage mapping methods and results.**

**File S2. Linkage map.** (a) High-density linkage map of a westslope cutthroat trout (WCT) and dot-plot comparison between the WCT to an (b) Arlee rainbow trout and (c) Swanson rainbow trout (version 1).

