## Supplementary material for "The genome assembly of the westslope cutthroat trout, *Oncorhynchus lewisi*, reveals interspecific chromosomal rearrangements with the rainbow trout": File S1

**SUPPLEMENTAL MATERIAL – LINKAGE MAPPING**

**METHODS**

**MAPPING CROSSES**

Diploid and haploid families used for linkage mapping were created using gametes collected from mature Westslope Cutthroat Trout (WCT) originating from the Danaher and Youngs Creek captive hatchery populations. These populations are housed at Sekokini Springs Hatchery (Columbia Falls, Montana), operated by Montana Fish, Wildlife, and Parks (MTFWP). Danaher and Youngs Creeks are located in the headwaters of the South Fork of the Flathead River drainage, upstream of Hungry Horse Dam. Routine genetic analyses conducted by MTFWP have repeatedly indicated that these populations are free from non-native Rainbow Trout and Yellowstone Cutthroat alleles (Ryan Kovach, personal communication).

Gynogenetic haploid families were created by pseudo-fertilizing eggs with irradiated sperm (Seeb & Seeb 1986; Limborg et al. 2016). Sperm was mixed 1:10 with milt extender solution and stored on ice immediately after extraction from the body cavity. The milt extender solution was prepared by mixing 8 parts Extender 8 solution (4 mM KCl, 135.5 mM NaCl, 4.2 mM NaH2PO4·H2O, 1.25 mM MgSO4·7H2O) and 2 parts Solution 2a (0.5% NaHCO3, 0.5% glucose). Motility was verified by mixing a drop of sperm from each male with activator solution (9 g NaCl, 1.5 g Glycine, 1.2 g Tris base mixed with 1 L of ambient water) and inspecting the sample under a microscope. Samples from males with low motility were discarded. Sperm was irradiated in a lightproof aluminum box with a Phillips (Amsterdam, Netherlands) 15W germicidal UV light bulb (Model TUV 15W/G15 T8). Aliquots were treated in clear Pyrex baking dishes for 10 minutes at a light intensity of 120 W/mm2 and were mixed halfway through treatment. Treated sperm was mixed with activator solution before being used to fertilize eggs and motility was again checked using the microscope. Eggs were incubated until the eyed stage and all embryos were preserved in 95% ethanol.

**LABORATORY METHODS**

DNA was extracted from diploid and haploid offspring using an isopropanol precipitation method after manually separating embryos from yolk. DNA quality (260/280 ratio) and quantity were measured using a Nanodrop 2000 Spectrophotometer (Thermo Scientific, Waltham, Massachusetts). The concentration of double-stranded DNA (dsDNA) was measured using QuantIt Picogreen assays (Thermo Fisher Scientific, Waltham, Massachusetts). A subset of offspring from each of 6 families was genotyped at 96 loci using the Fluidigm EP1 platform to verify ploidy. Results confirmed that a subset of offspring from each family were haploid. The two haploid families with the highest proportion of haploid offspring, along with one diploid family, were selected for RAD sequencing.

SbfI RAD libraries were prepared using the BestRAD protocol (Ali et al. 2016) and NEB Next Ultra DNA library preparation kits (New England Biolabs, Ipswich, Massachusetts). RAD libraries were sheared to a target fragment size of approximately 350 base pairs using a Covaris E220 Ultrasonicator (Covaris Inc., Woburn, Massachusetts) after ligating individual-specific 8 base pair barcodes to the sticky ends produced by SbfI digestion. Libraries were amplified for 12 cycles using plate-specific indexing primers, purified using Ampure XP beads, and quantified in triplicate using Quantit Picogreen assays. Libraries were sequenced in the 2x150 paired-end format on an Illumina HiSeq X instrument (Illumina, San Diego, California).

**BIOINFORMATICS**

We assembled RAD loci and called genotypes using tools available in Stacks v2.55 (Rochette et al. 2019) in combination with other software. Read quality was assessed using FastQC (Andrews 2010), and a custom Perl script was used to exchange read 1 and read 2 whenever the sample-specific barcode was located at the beginning of read 2 rather than read 1. Reads were then demultiplexed by sample and quality filtered using process_radtags. Specifically, we removed reads containing uncalled bases (Ns), removed reads if they contained any 15bp sliding windows with a mean base quality less than 10, and rescued reads with cut site or barcode mismatches by specifying the minimum edit distance among all BestRAD barcode pairs (options= --paired, -c, -q, -w 0.1, -s 10, -t 140, --inline_null, -e sbfI, --rescue, --barcode_dist_1 2, --retain_header). We then removed PCR duplicates using clone_filter (default settings) and excluded reads containing adapter sequence using Trimmomatic v0.32 (Bolger et al. 2014; options = ILLUMINACLIP:adapters.fasta:2:30:10, MINLEN:140). We counted the number of remaining reads for each individual, evaluated the histogram of read counts, and removed samples for which the log10 read count was more than 2 standard deviations less than the mean.

RAD loci were identified by running ustacks on all individuals (-M 5 -m 4 -H --max-locus-stacks 4 --model-type bounded --bound-high 0.05 --disable-gapped), and a locus catalog was generated using output obtained for all parents using cstacks (options= -n 3, --disable-gapped). Alleles and loci for all individuals were then matched against this catalog using sstacks (--disable-gapped). Output was then converted to BAM format using tsv2bam. SNP and RAD haplotype genotypes were called using gstacks and the maruki_low genotyping model (Maruki and Lynch 2017), with a variant alpha threshold of 0.01 and a genotype alpha threshold of 0.01. Genotype calls were converted to VCF format using the populations module (--batch-size 20000 --min-mac 5 --filter-haplotype-wise -r 0.7 -R 0.7 --vcf). Haplotype calls produced by Stacks were used for linkage mapping.

Non-haploid offspring were identified by plotting the number of high-confidence heterozygous genotypes (genotype quality > 30) against the number of observed non-maternal alleles. This resulted in the exclusion of 5 individuals. For haploid families, we used the maximum likelihood approach described by Waples et al. (2016) to identify collapsed duplicated loci with mappable segregation patterns and determine paralog genotypes. All settings were consistent with those of Waples et al. (2016), except we allowed for up to 30% missing data at a locus within each family. Genotypes for collapsed paralogous loci with mappable segregation patterns and singleton loci were exported to MSTmap format, then converted to Lepmap3 format using scripts from Blumstein et al. (2021).

For the diploid family, loci were removed if genotypic distributions in at least one haploid family suggested the locus was a collapsed duplicate (according to methods described above). Loci were also removed if more than 4 alleles were observed across all individuals, if neither parent was heterozygous, if one or both parents had missing genotypes, or if genotypes for more than 3 offspring were unexpected given parental genotypes. The proportion of offspring genotypes that were inconsistent with parental genotypes was tabulated across loci and across individuals to identify and remove problematic individuals. Remaining unexpected offspring genotypes were recoded as missing data, given that these were likely the result of sequencing errors. Additionally, we tested for deviations from Mendelian expectations using a chi-square test for goodness of fit and removed loci with p-values that were significant after a false discovery rate correction (q = 0.1). Genotypes for remaining offspring were exported to LepMap3 format and combined with those from the two haploid families.

**LINKAGE MAPPING**

We used various programs distributed with LepMap3 (Rastas 2017) to cluster markers on linkage groups, order markers, and verify the quality of the final map. For haploid families, genotypes for the male parent were encoded as homozygotes (A/A), and female parent genotypes were encoded as heterozygotes (A/C). Genotypes for homozygous C/C offspring were encoded as A/C heterozygotes, while genotype encodings for homozygous A/A offspring were left unchanged. Initial marker clustering was conducted using SeparateChromosomes2, using only male-informative markers, a logarithm of odds (LOD) threshold of 3, and a minimum marker count of 50. Additional markers were merged to this framework map using JoinSingles2All with the LOD threshold and LOD difference set to 3. Additional markers were iteratively merged to the map until no additional markers could be added. Markers were ordered using OrderMarkers2 with the minError parameter set to 0.001, the recombination parameters for both sexes set to 10^-6^, numMergeIterations set to 1, and all other parameters set to defaults. We conducted 100 iterations of the ordering algorithm and selected the order that maximized the likelihood of the data for each linkage group. Centromere positions were determined using the RFm method (Limborg et al. 2016).

**RESULTS**

We identified 33 linkage groups and determined positions for 19,604 markers. Of these, 18,540 (94.6%) were singleton markers and 1,064 (5.4%) were confounded paralogs with mappable segregation in haploid families. 3,743 markers were only informative in the male parent that produced the diploid family and 13,491 were informative in female parents. A total of 6 markers were heterozygous in both parents in the diploid family with no alleles shared between parents. A total of 361 markers were heterozygous in both parents in the diploid family with one allele shared between parents. 2,003 markers were heterozygous in the diploid family with both parents having the same alleles. The total female map length was 2,561 centimorgans (cM) and the total male map length was ,1201 cM. Centromere mapping and the distribution of map lengths indicated that 19 linkage groups were metacentric or submetacentric and 14 were acrocentric or telocentric. We identified 16 chromosome arms with high densities of mapped duplicated loci.
